# High-accuracy prediction of 5-year progression-free melanoma survival using spatial proteomics and few-shot learning on primary tissue

**DOI:** 10.1101/2024.10.30.620850

**Authors:** Anna Möller, Christian Ostalecki, Anne Hartebrodt, Katharina Breininger, Michael Erdmann, Stefan Uderhardt, Andreas Baur, David B. Blumenthal

## Abstract

Emerging techniques in imaging-based spatial proteomics (ISP) enable in-depth insights into the architecture and protein abundance of tissue(s). Explainable machine learning (xML) models promise to yield substantial advances in ISP data-based diagnosis and prognosis. However, a clinical application of these new possibilities predicting the course of a tumor has not been suggested yet. Here, we use a few-shot learning workflow on histological multi-antigen images to predict 5-year progression-free survival (PFS) in melanoma. We address the problem of a relatively small cohort (*n* = 22), by utilizing a pre-trained convolutional neural network (CNN) model, which we further pre-train on a proxy task for which more samples were available (*n* = 39) before fine-tuning for PFS prediction. Our approach yielded a model achieving an accuracy of more than 90%, outperforming baseline models trained on clinical data by around 10%. Using an xML technique, we identified immune infiltration and proteins associated with tumor progression as crucial predictors. This indicated that our models’ PFS predictions are not only highly accurate but also grounded in a relevant biological background.

## INTRODUCTION

In the rapidly growing field of spatial ‘omics’, several imaging-based spatial proteomics (ISP) techniques have been established to capture protein abundance in tissue^1–5^. Since most ISP data generation platforms are still relatively young, ISP data is typically used in pre-clinical studies which aim to elucidate tissue architecture or disease biology. In many of such mechanistic and functional studies, machine learning (ML) methods, especially convolutional neural networks (CNNs), are used for tasks like cell segmentation, cell classification, and differential analysis of model systems^4,6–8^.

However, ML-based analysis of ISP data also holds great potential for clinical applications, in particular, in high-precision personalized diagnosis and prognosis. For a successful clinical application of ML-based ISP analysis, the models need to have two properties: interpretability and generalizability. Firstly, as ML models in general, and CNNs with several millions of parameters in particular, are often blackboxes with limited interpretability and thus low utility for clinical decision-making. To address this, explainable machine learning (xML) techniques can be used. These techniques allow extraction of image regions to which an ML model attributes its decision — so-called regions of interest (ROIs)^9^ — making the models interpretable to the clinical decision maker. Secondly, the low number of patients paired with the high dimensionality and high resolution of the ISP images leads to an increased risk of overfitting when training ML models. To mitigate this risk, few-shot learning approaches, where ML models are first pretrained on auxiliary tasks with higher data availability and are then fine-tuned towards the specific downstream task have been succesfully used to improve model generalizability^10–14^.

Here, we explored an explainable few-shot learning workflow for ISP data, focusing on the clinically highly relevant use case of predicting 5-year progression-free survival (PFS) from primary malignant melanoma (MM) samples. We rely on ISP data from primary fresh frozen tumor samples of 27 melanoma and 12 benign nevi patients, which we generated between 2018 and 2020 with a prototype of the multi-epitope ligand cartography (MELC) technology^1^ available at the University Hospital Erlangen (UKER). Since all patients were diagnosed and treated at UKER, we were able to collect long-term outcome data, enabling us to train 5-year PFS prediction models. Remarkably, our models achieved accuracies of more than 90%, outperforming baseline models trained on clinical data by around 10%. Moreover, thanks to the use of xML techniques, our workflow revealed that our models base their predictions on ROIs with differential abundance of macrophages, granulocytes and CD4 cells between patients with favourable and unfavourable outcome, as well as proteins including TP73, KRT14, involved in tumor progression and adaptive anti-tumor immunity.

## RESULTS

### Explainable few-shot learning workflow for ISP data

MELC is an advanced immunohistochemistry technique that allows for the detection of numerous protein targets on a single tissue sample. This method employs sequential rounds of antibody staining, imaging, and photobleaching to remove fluorescent signals. The process is fully automated, enabling the visualization of hundreds of different proteins in one sample. After each imaging cycle, the fluorescent signal is bleached, allowing for the next round of staining without interference from previous markers. Here, we used an extensive immunostaining panel (see Key Resource Table), covering various immune cell types (e.g., CD4+ or CD8+ T cells) but also molecular processes such as cell adhesion (e.g., CD44, CD56) or apoptosis (e.g., Bcl-2, CD95) and proliferation pathways (e.g., Ki67, EGFR).

Our main goal was to predict 5-year PFS from the resulting spatially resolved protein abundance in primary melanoma tissue. To address this goal despite the relatively small number of MM patients in our cohort for whom PFS information is available (8 patients with PFS *<* 5, 14 patients with PFS ≥ 5), we designed the explainable few-shot learning workflow shown in Figure 1 (see Methods for details). Our workflow is based on ResNet18, an off-the-shelf CNN. We downloaded publicly available weights obtained via pre-training on histo-pathological images for this network to incorporate domain knowledge^15^. Then, we additionally made use of ISP data for 5 MM patients without PFS information (resulting in 27 MM patients in total) and 12 benign nevus (BN) patients. We started by pre-training the first convolutional layer of our adapted ResNet18 CNN on the auxiliary task to distinguish MM from BN. Since the information of whether a lesion is a BN or a MM is available in the intended use case to predict PFS for already diagnosed MM patients, we could use all MELC images for this pre-training step and did not have keep a separate test set to avoid data leakage^16^. Like this, pre-training on the proxy tasks allows the CNN to learn initial representations of the ISP data that facilitate downstream PFS prediction.

**Figure 1.**
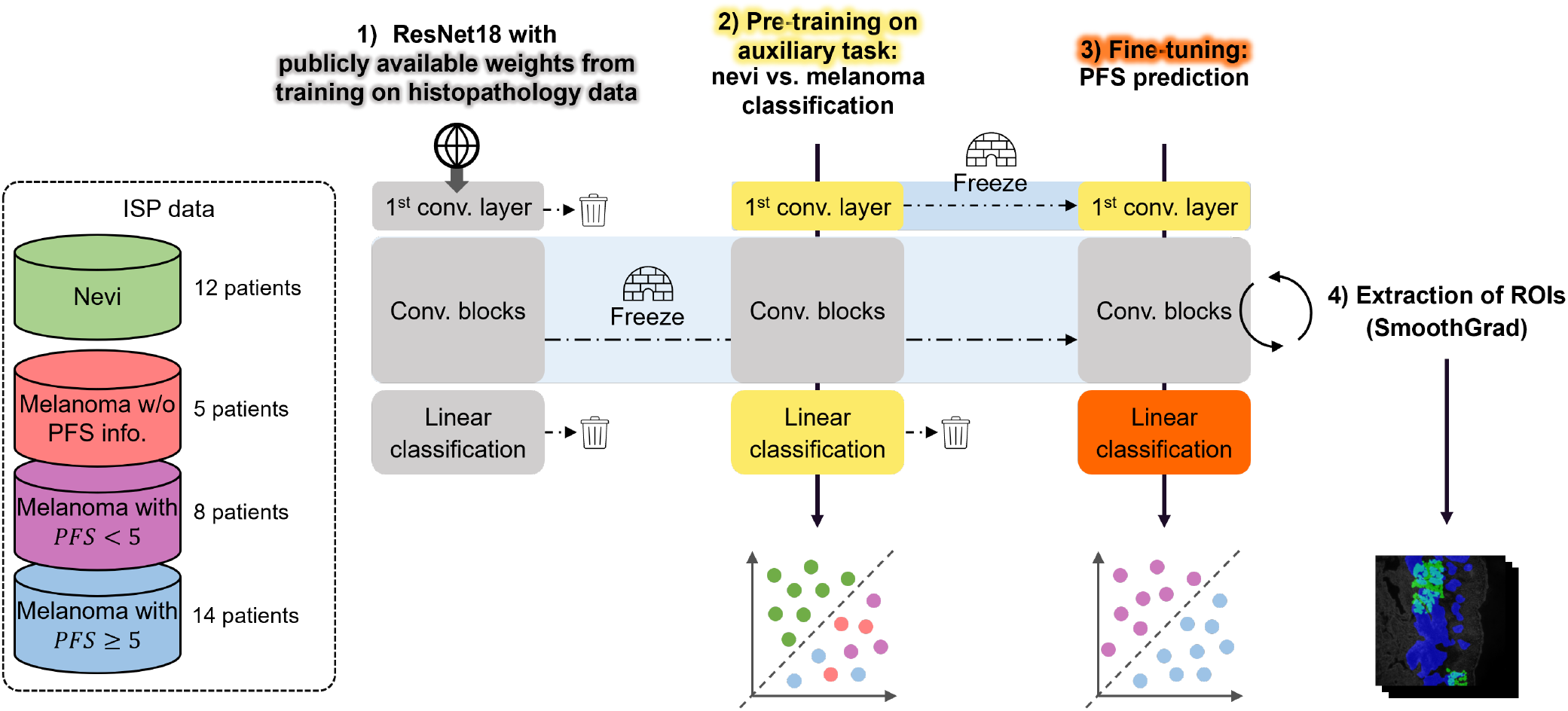
Schematic representation of the few-shot learning workflow. A neural network with weights trained on histopathology data (1) was further pre-trained to distinguish MM (red, purple, blue) from BN (green) in the high-dimensional ISP data (2). Then, the MM MELC images with missing PFS information (red) and the BN MELC images (green) were removed and the final layers of the model were trained for the actual task, the separation of melanoma MELC images with PFS < 5 (purple) from those with PFS ≥ 5 (blue) (3). The colors of the models’ constituents indicate for which task they were optimized. The models were validated via LOOCV and ROIs underlying the models’ decisions were extracted (4) and examined w. r. t. cell type composition and cell-level protein abundance differences between patients with PFS < 5 and PFS ≥ 5.

In a second step, we fine-tuned our pre-trained model to predict PFS, using the ISP data for MM patients for whom PSF information is available. To achieve realistic performance estimates, we here used leave-one-patient-out cross-validation (LOOCV) and computed performance metrics for MELC images from the patient in the test folds which were not used for fine-tuning. To explain the models’ decisions, we then extracted ROIs that were important to the decision, using the SmoothGrad^17^ feature attribution method, which identifies pixels that drive the CNN’s predictions. Finally, we analyzed the ROIs w. r. t. differences in cell type composition and cell-level protein abundance between MM patients with favourable and unfavourable outcome, thereby extracting biologically interpretable tissue characteristics that inform the models’ decisions.

### Deep learning models predict PFS with high accuracy

In order to decipher spatial architecture in primary tumor tissue, we carried out MELC imaging for our 27 MM and 12 BN patients. For each tumor sample, sections (each 1 mm^2^) of representative slides in the tumor area and surrounding microenvironment were analyzed by MELC. The number of analyzed sections depended on the size of the tumor. Each section was stained with 51 antibodies, each antibody giving one staining. In the following, we refer to all 51 stainings of a section as a (51-dimensional) “MELC image”. Figure 2a visualizes the numbers of MELC images for MM patients with PFS ≥ 5 and PFS < 5, the underlying numbers of patients and MELC images are summarized in Table 1. We computed performance metrics for our CNN models on two levels. While MELC image-wise results are the direct output of the CNN models (our CNN models yield predictions for MELC images and our dataset contains several MELC images per patient), patient-level predictions are more relevant from a clinical point of view. To obtain patient-level predictions, we averaged the MELC image-level scores produced by the CNN model for all MELC images of one patient. We compared our CNNs to two baseline models. The first baseline model is a random forest classifier (RFC) that was trained on six clinical features (sex, age, tumor thickness, tumor ulceration status, coarse tumor location). The RFCs were trained and evaluated with the above-mentioned LOOCV setup also used for our CNNs. To address data imbalance on the different labels, the second baseline is are biased random classifier that randomly predicts the outcome following the prior probabilities of the ground truth labels on MELC image and patient level.

**Table 1:** Numbers of patients and MELC images stratified by condition and PFS status. For the statistical analyses of the cell type fractions and protein abundances, we discarded MELC images with an unsatisfactory cell segmentation result.

| Condition | PFS status | Patients | MELC images | MELC images used for analysis |
| --- | --- | --- | --- | --- |
| Melanoma | PFS ≥ 5 | 14 | 97 | 76 |
| Melanoma | PFS $<$ 5 | 8 | 15 | 14 |
| Melanoma | N/A | 5 | 13 | — |
| Benign nevi | — | 12 | 57 | — |

**Figure 2.**
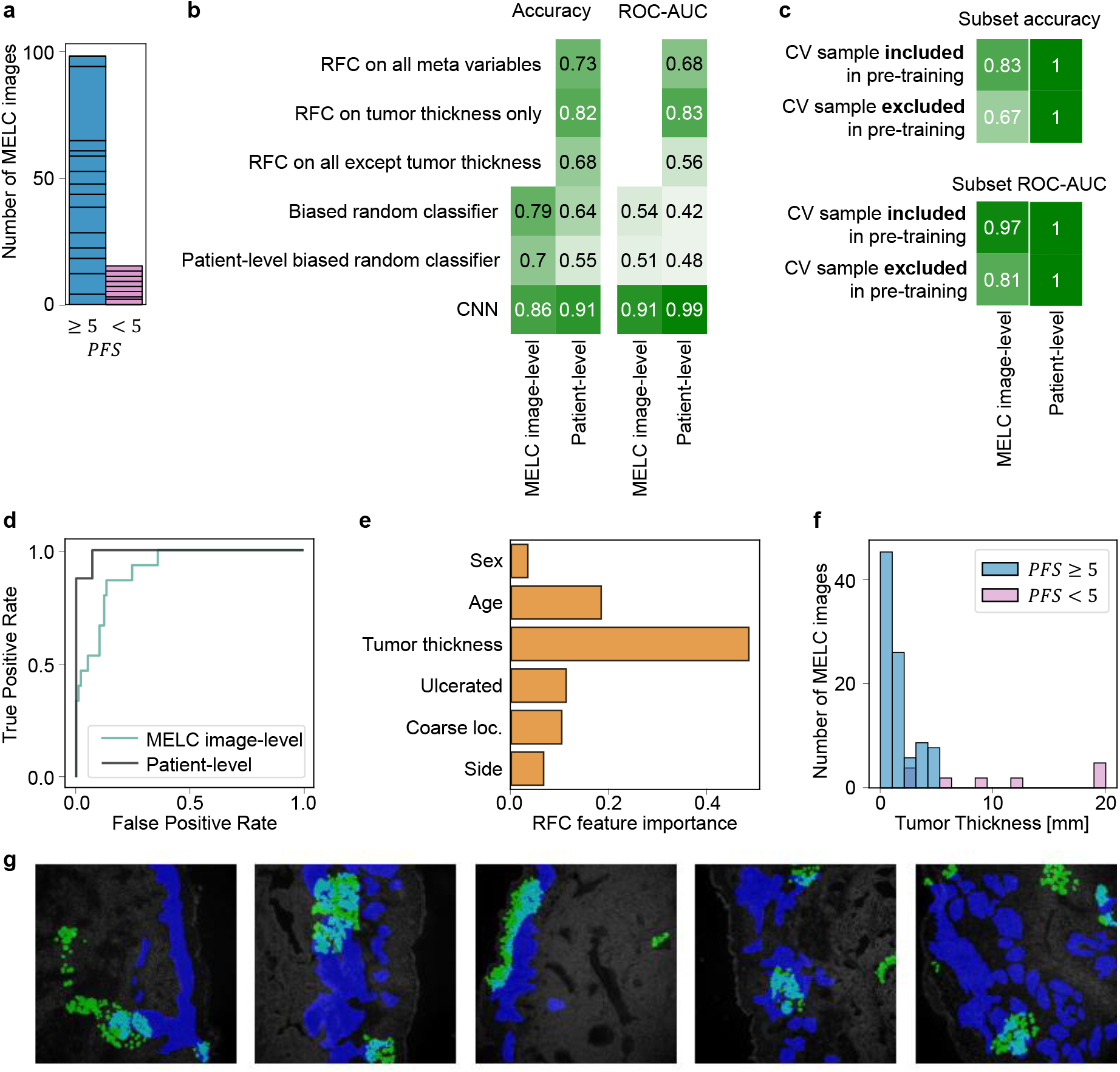
Performances of the ML classifiers. **a** Number of MELC images by PFS group, individual patients are indicated by boxes. **b** LOOCV accuracies and ROC-AUC scores obtained by the tested ML models on the different levels. PFS < 5 is encoded as 1, PFS ≥ 5 as 0. Scores are binarized by assigning all MELC images and patients with (average) prediction *ŷ* ≥ 0.5 to the positive class. Accuracies are calculated based on the binarized scores. Further metrics are reported in Table S1, age- and sex-resolved results are shown in Figure S1. **c** Accuracies and ROC-AUC scores for a randomly chosen subset of four patients remain high even when the corresponding MELC images are completely removed from the pre-training. Further metrics and sex and age resolved performance are reported in Table S2. **d** The ROCs for the CNN’s predicted probabilities on the different levels show excellent separability. **e** Averaged impurity-based feature importances of the cross-validation RFCs. **f** Tumor thickness distributions in tissue sections from MM patients with PFS≥ 5 and PFS < 5. **g** Overlay of segmented cells in the PFS model’s ROIs (green), expert-marked tumor regions (blue), and their intersection (cyan) on the phase image.

The RFCs based on clinical features obtained an accuracy on the patient level of 0.73, taking all available variables into account. Consistent with previous studies^18^, the best performance of the RFC models was obtained when using only tumor thickness as feature (patient-level accuracy of 0.82), whereas removing the information on tumor thickness leads to a drop in performance to 0.68 (Figure 2b). Investigating the feature importance distributions of the cross-validated RFCs (Figure 2e) and the tumor thickness distributions in MM patients split by 5-year PFS status (Figure 2f) further suggests that tumor thickness is a strong indicator for PFS. While the primary purpose of including RFC trained on clinical data was to benchmark against our CNN models, these results are also relevant in themselves: To the best of our knowledge, there are no prior studies where ML models are trained to predict 5-year PFS from clinical melanoma data.

Our CNN models further improved upon the performance of the RFCs trained on clinical data and the biased random labels, and achieved a patient-level accuracy of 0.91 and a patient-level ROC-AUC of 0.99 (Figure 2b, Figure 2d). To assess the impact of pre-training on the proxy task of MM vs. BN classification, we randomly selected 4 patients from our MM cohort (2 of each class) and compared the LOOCV performance metrics obtained when including and excluding the corresponding MELC images during pre-training (we restricted this analysis to 4 patients since it requires time-consuming repeated pre-training for each patient). Interestingly, performances dropped on the MELC image-level, but not on the patient-level (Figure 2c). On the one hand, this result shows that pre-training on the proxy task indeed helps the CNNs to learn ISP data representations useful for PFS prediction. On the other hand, it indicates that in an application scenario where PFS should be predicted for a new patient and repeated pre-training is practically infeasible (e. g., due to limited access to hardware, ML expertise, or data at the point of care), an adapted workflow that does not require repeated pre-training would yield similarly accurate PFS prognoses. This shows that, depending on the availability of hardware, data, and ML expertise, different configurations of our xML workflow could be used in a clinical setting.

In order to characterize the ROIs used by our model, we manually annotated tumor regions in a subset of 5 MELC images from our MM cohort with PFS information. Figure 2g shows the model-extracted ROIs for these images (green) in comparison to the expert-annotated tumor regions (blue). A substantial fraction (roughly 52%) of the cells in the ROIs were not contained in the tumor regions, showing that our prediction models not only relied on information from the tumor itself but also from the micro-environment.

### Explainable ML reveals relevance of immune infiltration for PFS

To further elucidate which information was used by our CNNs, we calculated cell type fractions within the ROIs for each MELC image. To identify cell types, we transferred cell type annotations from healthy skin tissue single-cell RNA-sequencing (scRNA-seq) data obtained from GTEx, using bimodal distribution fitting (see Methods for details). Figure 3a shows the obtained MELC image-level distributions, split by PFS status. The most pronounced differences between melanoma patients with favorable and unfavorable outcome can be observed for granulocytes, macrophages, and suprabasal keratinocytes. Especially the changes in granulocyte and macrophage density are interesting in the light of existing studies: A potential reason for the higher fraction of granulocytes in the cohort with PFS ≥ 5 could be the presence of eosinophilic granulocytes in the tumor micro-environment, which have been associated with a favorable outcome in MM and other solid tumors^21–23^. Conversely, increased macrophage density has previously been reported to be correlated with tumor thickness^24^, which Antohe *et al*.^18^ identified as a strong predictor for PFS. This is well aligned with the substantially higher fraction of macrophages in ROIs of MELC images from patients with PFS < 5 (Figure 2f). Generally, macrophages and melanoma tumor cells show complex interactions in opposing directions, activating the immune response but simultaneously promoting growth^25,26^. Our results point to a less favorable outcome for patients with a higher macrophage density.

**Figure 3.**
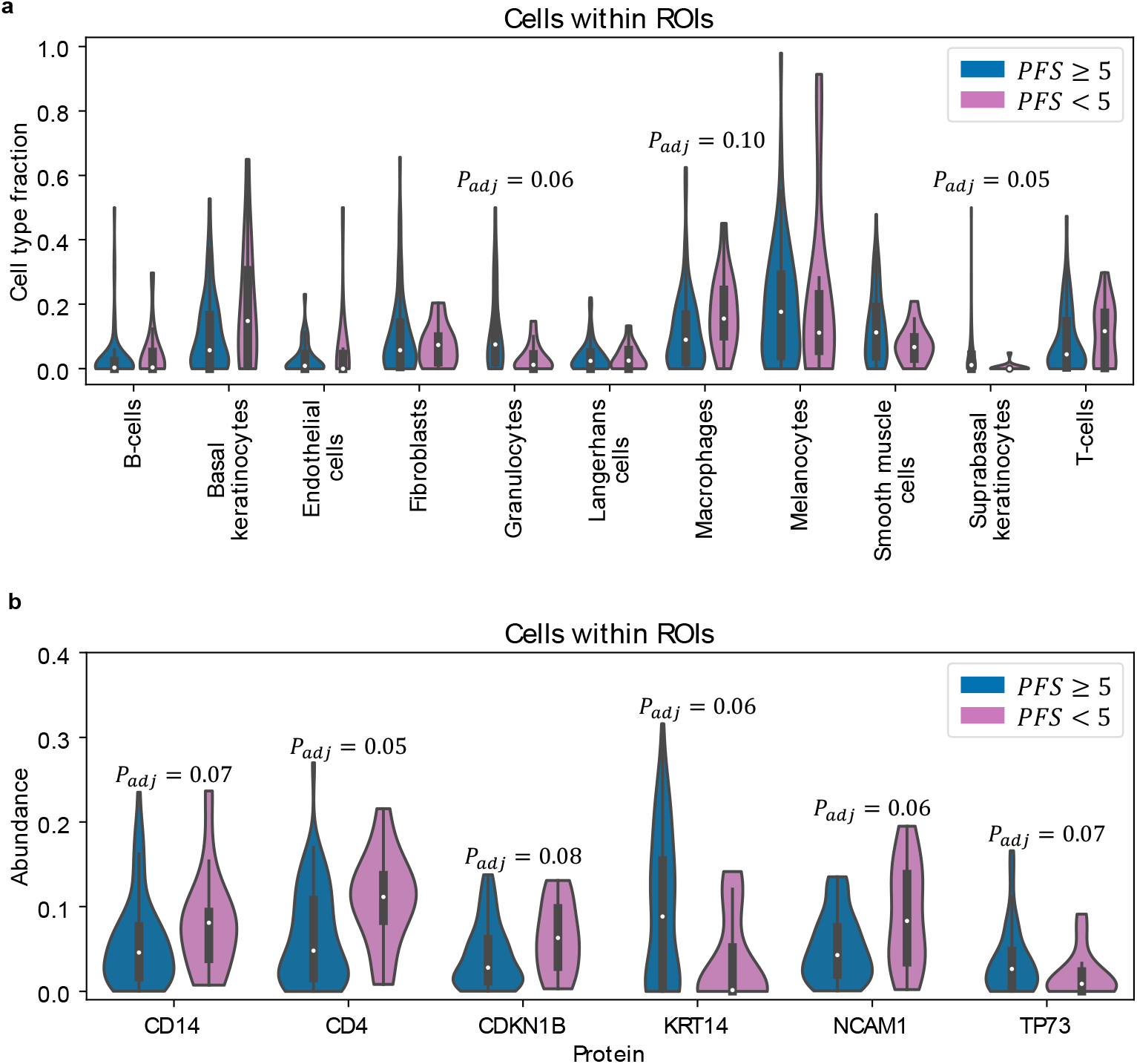
Differences in distributions of cell type fractions (**a**) and protein abundances (**b**) within ROIs of MELC images from MM patients with PFS ≥ 5 and PFS < 5. *P* -values were computed with the two-sided Mann-Whitney U test^19^. Benjamini-Hochberg correction^20^ was used to correct for multiple testing and to adjust the false discovery rate (FDR). Adjusted *P* -values with *P*_*adj*_ < 0.1 are shown.

Similarly to the cell type fractions, we also determined the average abundances of different proteins within the ROIs of each MELC image and again compared the obtained distributions between MM patients with PFS ≥ 5 and PFS < 5. Figure 3b shows the results for the proteins for which we obtained the largest differences. All of these proteins have been reported to play a role in MM^27–41^ (see Table S3 for a detailed discussion of the individual markers).

Although many of the observed differences in cell type densities and protein abundancies within the ROIs extracted by our models do not reach statistical significance after multiple testing correction, we hence observe a strong enrichment in cell types and proteins that have previously been linked to MM progression. On the one hand, this provides substantial evidence that our CNN models indeed pick up on true biological signal in the ISP data to predict PFS. On the other hand, the mostly weak associations of individual proteins and cell types with PFS also suggests that ML models such as our CNNs are needed to aggregate weak individual signals, highlighting their potential for high-precision personalized prognosis in MM care.

## DISCUSSION

In this study, we successfully trained CNN models to accurately predict 5-year PFS from primary tumor ISP data in a few-shot learning setup. By pre-training our models on a proxy task (BN vs. MM classification) for which additional MELC images were available, we obtained ISP data representations that allowed excellent separation between MM patients with favourable and unfavourable long-term outcome.

While imaging-based ML models have been used extensively in MM research and have yielded impressive results especially for diagnostic tasks^42,43^, our study is one of the first attempts to predict long-term outcome from primary tumor ISP data. A closely related work is the study by Meiser *et al*.^8^, who used few-shot xML for ISP data to identify characteristic clustering behaviour of dendritic cells and T cells as a potential explanation for differences in protective immunity between COX-deficient and control BRAF^V600E^ mouse models. Inspired by the work of Meiser *et al*., we showed that few-shot xML for ISP data is not only a valuable tool for the analysis of data from model systems, but can also be used for explainable, high-accuracy outcome prediction, potentially paving the way for personalized patient monitoring in MM care. Another related work is the study by Van Kleunen *et al*.^44^ who used graph-based representations of ISP data generated via multiplexed ion beam imaging^4^ to predict overall survival and PFS in ovarian cancer. Despite a substantially larger cohort (*n* = 83), their best-performing models only achieved an AUC of 0.71 (much lower than our models), indicating that pre-trained imaging-based models as employed here may be more promising for clinically relevant predictions based on ISP data.

Our workflow includes ROI extraction coupled with image segmentation and cell type assignment to facilitate the analysis of the tissue regions that most contribute to the models’ decisions. We identified a significantly higher cell type fraction of macrophages and a lower fraction of granulocytes in the ROI cells of patients with poor outcome compared to the other group. Furthermore, comparing the protein abundances in the ROI cells of the two groups, we observed the most significant expression differences for proteins (CD4, CD14, CDKN1B, KRT14, NCAM1, and TP73) with strong literature support for association with cancer progression. These results add explicability to our models’ predictions, which is a key component of the EU’s ethics guidelines in trustworthy AI^45^ and is hence crucial for a potential translation into clinical care^46^.

Independently of the proposed xML workflow, the annotated MELC dataset is itself a very valuable resource, as only few ISP datasets from tumor patients with similarly rich clinical annotations including information of 5-year PFS exist. To facilitate further advances in ISP-based ML research for diagnosis and prognosis, we published the anonymized ISP and clinical data along-side this article (https://doi.org/10.5281/zenodo.10996361).

Our study opens up several avenues for future work. Firstly, we analyzed ROI cells only w. r. t. differences in cell type composition and protein abundance. In addition to these basic analyses, our data allow numerous other possibilities to analyze the ROI cells, e. g., using tools such as GraphCompass^47^ or SHouT^48^ that quantify differences in spatial tissue organization between two conditions. Running such tools on the identified ROIs may reveal that the predictions of our CNN models are also informed by more complex tissue patterns such as clustering behaviour of specific cells, which could further improve the explicability of the models’ predictions.

Secondly, the proposed explainable few-shot learning workflow can be adapted to other supervised classification or regression tasks *T* that are based on ISP data. The only prerequisite is that additional ISP data are available to allow fitting the weights of the initial convolutional layer on a proxy task which is sufficiently similar to real prediction task *T* . Since data preparation for supervised ML model training almost always leads to the exclusion of some samples due to lacking or incomplete annotations, defining such a proxy task is likely often possible. The workflow proposed here could therefore help fitting ML models on ISP datasets with small cohort sizes in a wide variety of applications.

## Limitations of the study

Challenges arise from some of the dataset’s properties: Some of the images could not be used for cell-level analyses due to staining artifacts which limited the spatial segmentations. Staining artifacts occur when tissue is not optimally frozen (despite applying an automated procedure), often due to the nature of the excised tumor (e. g., ulcerated versus not ulcerated). This reduced the number of available MELC images, but did not affect the overall results. Moreover, the ISP data was generated with a prototype of the MELC imaging platform and we hence had to rely on our own prototype software to pre-process the raw MELC images. While we are confident that our pre-processing methods (cell segmentation, protein abundance quantification, cell type calling) are adequate for out data (see Methods for detailed justifications), different methodological choices in the individual pre-processing steps might have led to slightly different results.

Another limitation is introduced by the general architecture of CNN models and the fact that SmoothGrad is applied to the last layer of the model to extract the ROIs. This measure is reasonable, as the last layer is regarded to contain the most abstract spatial information on the input image, but it comes with the disadvantage that the receptive field that produces the spatial features is rather coarse. In our case, the output of the last convolutional layer of ResNet18 is of size 16 *×* 16, so this is the most fine-grained resolution the extracted ROIs can possibly have. This implies that the edges of the extracted ROIs are somewhat blurry, which may have led to small distortions in our differential cell type fraction and protein abundance analyses. Feature attribution methods in general have undergone criticism of being unreliable, fragile, and sensitive to small perturbations^9,49–51^. However, SmoothGrad counteracts most of these problems, which is why we chose to use it for our study.

Finally, it is important to stress that, despite the inclusion of xML techniques, the PFS predictions of our CNN models are still not fully transparent and explicable to a human decision maker. Our results on differential cell type compositions and protein abundances constitute *post hoc* explanations of the models’ predictions, but ultimately, the decision boundaries are fit in high-dimensional embedding spaces that are impossible to directly inspect by humans. In view of this inherent limitation of deep learning models such as CNNs, incorporation of uncertainty quantification methods^52^ that allow to annotate model predictions with confidence scores could further strengthen the translational potential of the proposed few-shot learning workflow.

## Supplemental information index

Document S1: Tables S1 to S3 and Figures S1 to S4.

## Acknowledgments

A.M. and D.B.B. were funded by the Deutsche Forschungsgemeinschaft (DFG, German Research Foundation) – 516188180. A.M., A.H., and D.B.B. were supported by the German Federal Ministry of Education and Research (BMBF, grant no. 031L0309A). SU is supported by DFG (project-IDs 448121430, 405969122, 505539112), the Hightech Agenda Bavaria and by an ERC starting grant (NEXUS; project-ID 101039438).

## Author contributions

A.M. and D.B.B. conceived and designed this study and drafted the manuscript. A.M. preprocessed the data, developed and implemented the machine learning models, and analysed the results. C.O. generated the data. C.O. and M.E. annotated the data. K.B. and A.H. advised on the model training and evaluation. A.H. advised on the statistical analyses. S.U. provided biomedical expertise. D.B.B. and A.B. supervised this work. All authors reviewed and approved the manuscript.

## STAR METHODS

### Key resources table

Document Key Resources Table.

### Resource availability

#### Lead contact

Requests for further information and resources should be directed to and will be fulfilled by the lead contact, David B. Blumenthal.

#### Materials availability

This study did not generate new materials.

#### Data and code availability

The data underlying this study are available in anonymyzed form at https://doi.org/10.5281/zenodo.10996361. Source code to reproduce the reported results is available at https://github.com/bionetslab/xml_melc_melanoma/. An AIMe^53^ report with further details on data, models, and validation strategy is available at https://aime-registry.org/report/R9lTBD.

### Experimental model and study participant details

#### Study participants, ethics, collection of clinical data, and marker panel selection

All data underlying this study were generated from skin samples collected from adult MM and BN patients treated at UKER between 2010 and 2020. Numbers of patients and MELC images are summarized Table 1, age and sex distributions are reported in Figure S1. The ethnicities of the patients mirror the average population in northern Bavaria, Germany. The study has been approved by the Ethics Committee of the Medical Faculty of the Friedrich-Alexander-Universitä t Erlangen-Nü rnberg (approval date: July 5, 2023; approval number: 23-132-B).

Clinical and outcome data were curated by a dermato-oncologist at UKER based on in-house clinical records and data from the Bavarian Cancer Registry. All MM patients who survived at least 5 years without recurrence after presenting their primary tumor were assigned to the PFS ≥ 5 group. Patients who died of MM or had a recurrence within 5 years were assigned to the PFS < 5 group. For the remaining MM patients (lost follow-up, death of unknown cause, recurrence-free but with primary diagnosis less than 5 years ago), we set PFS = N/A. To determine the marker panel for MELC imaging, we screened over 500 antibodies on MM and BN tissue to assemble a set of antibodies with strong and specific staining patterns. This unbiased approach yielded 51 antibodies which produced high-quality images for the majority of the samples (Table S3).

#### Influence of age and sex on ML model performance

Figure S1b shows LOOCV performance metrics of our CNN models and the RFC baselines upon splitting the MM patients with PFS information by sex or median age. We observe that our CNNs yield similar accuracies for male and female patients, that they yield better predictions for younger than for older patients, and that the RFCs trained on clinical data are much more accurate for older than for younger patients and perform slightly better for women than for men. While this suggests that an ensemble approach which uses RFCs trained on clinical data for older patients and CNNs trained on ISP data for younger patients may be a promising option, the small sample sizes in the individual demographic subgroups make it impossible to draw strong conclusions from these observations.

### Method details

#### Multi-epitope ligand cartography (MELC) data generation

MELC imaging^54^ is a robot-automated whole-cell imaging technique that enables multi-antigen analysis in cells or tissue. For each surface marker of interest, the following process is repeated: First, fluorochrome-coupled antibodies (ligands) label specific antigens (epitopes) in the tissue section. Then, unbound antibodies are washed off, and the fluorescent image is captured using a microscope. The fluorochrome is deactivated through photobleaching so that only the signal from the last antibody-coupled fluorochrome remains in each image. The resulting fluorescence signals from each of the individual antibodies are then overlaid to create a tissue map of different epitopes. The MELC images underlying this study have been recorded at UKER from 2018 to 2020. The original resolution is 2018 *×* 2018 pixels for at least 50 markers, resulting in over 200 million pixel values per image. As input channels, we choose all marker images that have been recorded for the majority of samples, leading to 51-dimensional input images. The list of markers used for training the models can be found in Table S3.

#### Cell segmentation

We used StarDist segementation^55^, as provided by the StarDist Python library, on the bleach image of the Propidium iodide channel to segment the nuclei. As not only the nuclei, but also the cell membranes outlining the entire cells are of interest, we additionally applied StarDist segmentation to the CD45 (a transmembrane protein) image. We then overlayed nucleus and membrane signals and performed spatial mapping. We use these mappings to calculate an average relative membrane radius with regard to the nuclei size. For all nuclei where we did obtain a corresponding membrane signal, we drew a radius with the respective size around the nucleus center. If two circles overlapped, we assigned each pixel in the overlapping area to the closest nucleus center by calculating a nearest neighbor segmentation.

#### Cell-level protein abundance quantification

Due to the varying quality of MELC images, there is no unique mapping function from intensity to protein abundance across all images. To binarize the stainings images and account for noise and intra-image brightness variations, we used adaptive thresholding as provided by the OpenCV Python library. Each pixel is set to 1 if it is above the Gaussian-weighted mean of its vicinity plus a constant *C*. This calculation requires two hyper-parameters: the window size (*s, s*) and *C*. Qualitatively evaluating the results of a parameter search with the help of a biochemistry expert, the window size was chosen as (201, 201) and *C* was set to the standard deviation of the current sample. With this approach, we were able to correct salt-and-pepper noise as well as intensity distortion. We also tested other commonly used techniques like Otsu’s thresholding^56^, but found that the adaptive approach described above yields more robust results. Using the binary pixelwise expression, we calculate floating point expression per segment (e. g, cell) by dividing the number of active pixels (value 1) by the total number of pixels per segment.

#### Rule-based cell type assignment

Standard cell type assignment methods such as clustering followed by retrieval of cluster markers via differential expression analysis did not yield good results on our data. The main reason for this is that established cell type calling methods have primarily been designed for single-cell RNA sequencing (scRNA-seq) data with whole genome readout, whereas our ISP data only contained 51 protein channels. In view of this, we used a simple rule-base cell type assignment workflow, relying on bimodal distribution fitting and healthy skin scRNA-seq data with pre-computed cell type annotations from the Human Protein Atlas (HPA)^57,58^ as reference. Based on the HPA skin data, we identified the most characteristic markers for cell types annotated in the HPA data and constructed a list of marker-cell type pairs. We sorted this list by fold change, such that marker-cell type pairs at the top of the list contain the most characteristic markers for the corresponding cell type. Then, for each MELC image in our ISP dataset, we iterated through this list from top to bottom. For each marker-cell type pair, we tested if the cell-level protein abundance of the marker follows a bimodal distribution in the currently processed image. If so, we calculated a cutoff based on the means and standard deviations of the fit bimodal distribution and assigned the cell type to all cells with marker abundance above the cutoff. Then, we continued with the next marker-cell type pair. The different lists of marker-cell type pairs used for the assignment are shown in Figure S2.

#### Image data preparation and data augmentation for ML model training

For training and validation of ML models, the images were down-sized to 512 *×*512 pixels. We performed channel-wise normalization to zero-mean and unit-variance with statistics calculated on the dataset. To increase sample sizes, we moreover carried out content-invariant data augmentation, meaning that the input images were randomly rotated up to 30 degrees, resized with a scaling factor *s* ∈ [0.9, 1], or flipped horizontally or vertically. Furthermore, we randomly added Gaussian noise to single channels.

#### Pre-training of deep learning model

To minimize the risk of overfitting, we used a ResNet18 architecture with pre-trained weights^15,59^. We chose this architecture because it has a comparably small amount of parameters and good performance in natural image classification. We froze all parameters of the model except for the final classifier and the first convolutional layer, which needs to be replaced according to the input dimensions.

As the MELC images used for this study have *c* = 51 channels, we had to replace the first convolutional layer of the feature extraction part of the ResNet18 model, which originally expects three channels. Hence, we fit the parameters of this first convolutional layer on an auxiliary task, the classification of MM (encoded by label 1) versus BN (encoded by label 0) MELC images. To retrieve the hyper-parameters for pre-training this initial layer, we used patient-level crossvalidation. Specifically, we grouped the MELC images by patient. Each BN patient was combined with the MM patient with the most similar number of images as a validation set and was held out for one training run. We balance both the training and the validation set using the data augmentation techniques described above and optimized the first convolutional layer and the final classifier with regard to binary cross entropy loss, using the ADAM optimizer^60^.

We averaged the performance metrics of the different runs to retrieve the optimal learning rate and the number of epochs to be trained for. After 15 epochs with a learning rate *η* = 0.0005, we achieve a patient-level cross-validation F1-score of 0.81 and an accuracy of 0.8 on the individually balanced validation sets. The optimization processes and evaluation details of the cross-validation runs are shown in Figure S3. Subsequently, we used the entire dataset as training data for one final run with these hyper-parameters. Like this, the CNNs could learn meaningful representations for all MM MELC images to facilitate downstream PFS prediction.

We would again like to stress that this does not constitute data leakage, since we expect the information if a sample is a MM or a BN sample to be available when predicting 5-year PFS of already diagnosed MM patients at inference time. However, the proposed workflow does require repeated pre-training and availability of the entire dataset at inference time, which may be practically infeasible at the point of care. To investigate how much performance drops in a workflow that does not require pre-training at inference time, we therefore randomly selected 4 patients (two with PFS < 5 and two with PFS *>* 5) and additionally carried out both the pretraining phase and the downstream fine-tuning (see next paragraph) without the corresponding MELC images (see results reported in Figure 2c and Table S2).

#### Fine-tuning of deep learning models

We froze the weights of the pre-trained model and removed the linear classifier that was used to separate MM from BN after the convolutional blocks. On the truncated model, we appended multiple linear layers with dropout and ReLU, similar to VGG16’s last linear layers^61^. The feature extraction part of the model hence remains identical; we only appended and trained a new classifier. For this, we use LOOCV and performed *n* = 22 training runs. In each training run, all MELC images of one patient were held-out and we trained on the remaining 21 patients. Thereby, we avoided that the model learns to recognize the patients, instead of finding generalizable patterns in the samples that separate the conditions. For each run, we computed the model’s MELC image-level predictions for the patient not seen during training. We averaged the performance over the training runs and found that training the last fully connected layers with a learning rate *η* = 0.001 for only 5 epochs leads to the lowest validation loss. We initialized the parameters with Xavier initialization^62^ and minimized the binary cross entropy loss with ADAM optimizer^60^. Details on the optimization processes (training and validation loss, validation F1-score and accuracy, and learning rates over epochs) are shown in Figure S4.

#### Training of the baseline models

The dataset includes six section-level clinical variables encoded as binary or continuous variables for which data was collected when the patients presented at the clinic with the primary tumor: sex, age, tumor thickness, whether the tumor is ulcerated, and its coarse location. For the 22 melanoma patients with available PFS information, we first aimed to predict PFS from the clinical variables in a LOOCV setup with a RFC. For this, the clinical data of one patient was held-out and the remaining 21 patients were balanced by over-sampling the minority class and then used to train a RFC. Using the trained RFC model, we then predicted 5-year PFS status for the held-out MELC images. We repeated this procedure for all *n* = 22 patients and calculated the performance metrics based on the actual outcome data and the predictions. To implement the RFCs, we used the RandomForestClassifier class of the scikit-learn Python library with classweight=“balanced” and all other hyper-parameters left at defaults. For the biased random predictions, we randomly shuffled the labels on MELC image and patient level.

#### Extraction of regions of interest

The general idea of SmoothGrad^17^ is based on GradCAM^63^. It involves passing each image through the network, extracting the activation of the last convolutional layer in the forward pass, and multiplying it with the gradient of the same layer’s parameters with regard to the target label retrieved in the backward pass. For robustness and smoother results, this procedure is repeated *n* times (we used *n* = 10), and Gaussian noise is added to each image. For each noisy image, the result of the procedure is averaged channel-wise (to get rid of the depth dimension) and all negative values are set to zero. We corrected for artefacts by repeating this procedure with an all-zero image, hence, passing only Gaussian noise through the network, and subtracted this correction image from the actual feature attribution map. Then, the result was up-scaled to match the input image size. Averaging the feature attribution maps of the 10 noisy images yielded the final ROIs, representing which image regions have been important to the models’ decisions.

In the original publication, the concept is explained for multi-class scenarios, where the target label is back-propagated as a one-hot-encoded vector. As we only had a single class, and therefore scalar output, we negated the gradients if the target label is *y* = 0. In the original GradCAM publication^63^, this is advised for retrieving counter-factual examples for one of the multiple classes, which is what is required in a binary scenario for the samples with *y* = 0.

For each cross-validated model, we extracted the ROIs for the MELC images that were heldout, implying that they were not seen in the fine-tuning process for the parameters which play a role in the ROI extraction. The output of the SmoothGrad procedure is a floating point attribution map. For binary ROIs, we used the top decile of positive values in the result. We investigated how the SmoothGrad result output differed depending on which layer is taken into account. In the first layers, more fine-grained details (resembling edges or gradients) were highlighted almost everywhere in the image, whereas using the last layer yielded more selective areas similar to the medical expert’s tumor regions.

### Quantification and statistical analysis

Quantification and statistical analyses were carried out as follows:

- The exact number of patients and MELC images used in our analysis are shown in Table 1. We used the Python packages Pandas and Seaborn for data management and visualization.
- For cell type assignment, we assumed that the abundance of a marker protein *m* for a cell type *t* with a high-quality signal should follow a bimodal distribution across the cells of the stained image, where the two modes represent the cells which do or do not belong to cell type *t*. When processing a potential marker-cell type pair (*m, t*) in our iterative cell type assignment approach, we therefore fit a Gaussian mixture model with two components to *m*’s abundance distribution over the cells with yet undetermined cell type. Then, we used the statistics of the resulting components to determine the lower and the higher means *µ*_1_ and *µ*_2_ and the corresponding standard deviations *σ*_1_ and *σ*_2_. Subsequently, we checked if

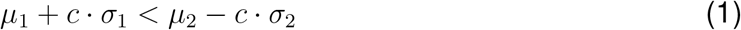

and if the Aikake information criterion (AIC) for the Gaussian mixture model was smaller than the AIC of a univariate Gaussian fit on the same data. If both conditions held, we assumed that we observed a bimodal distribution and assigned cells where *m*’s abundance was larger than *µ*_2_ *c σ*_2_ to cell type *t*. We chose *c* = 0.5 and validated this approach with careful examination of the results with the help of a medical expert. We used the GaussianMixture class from scikit-learn to fit Gaussian mixture models and compute AICs.

- All CNN models were trained and evaluated in the PyTorch framework.
- In the SmoothGrad procedure, we used Gaussian noise with 𝒩 (0, 1) with PyTorch’s randn function.
- To test for differences between cell type fractions and protein abundances, we used the mannwhitneyu function from the SciPy Python package for testing and the multitest from the statsmodels Python package for *P* -value adjustment.

