## Supplement for "High-accuracy prediction of 5-year progression-free melanoma survival using spatial proteomics and few-shot learning on primary tissue"

### 1 Supplementary Figures

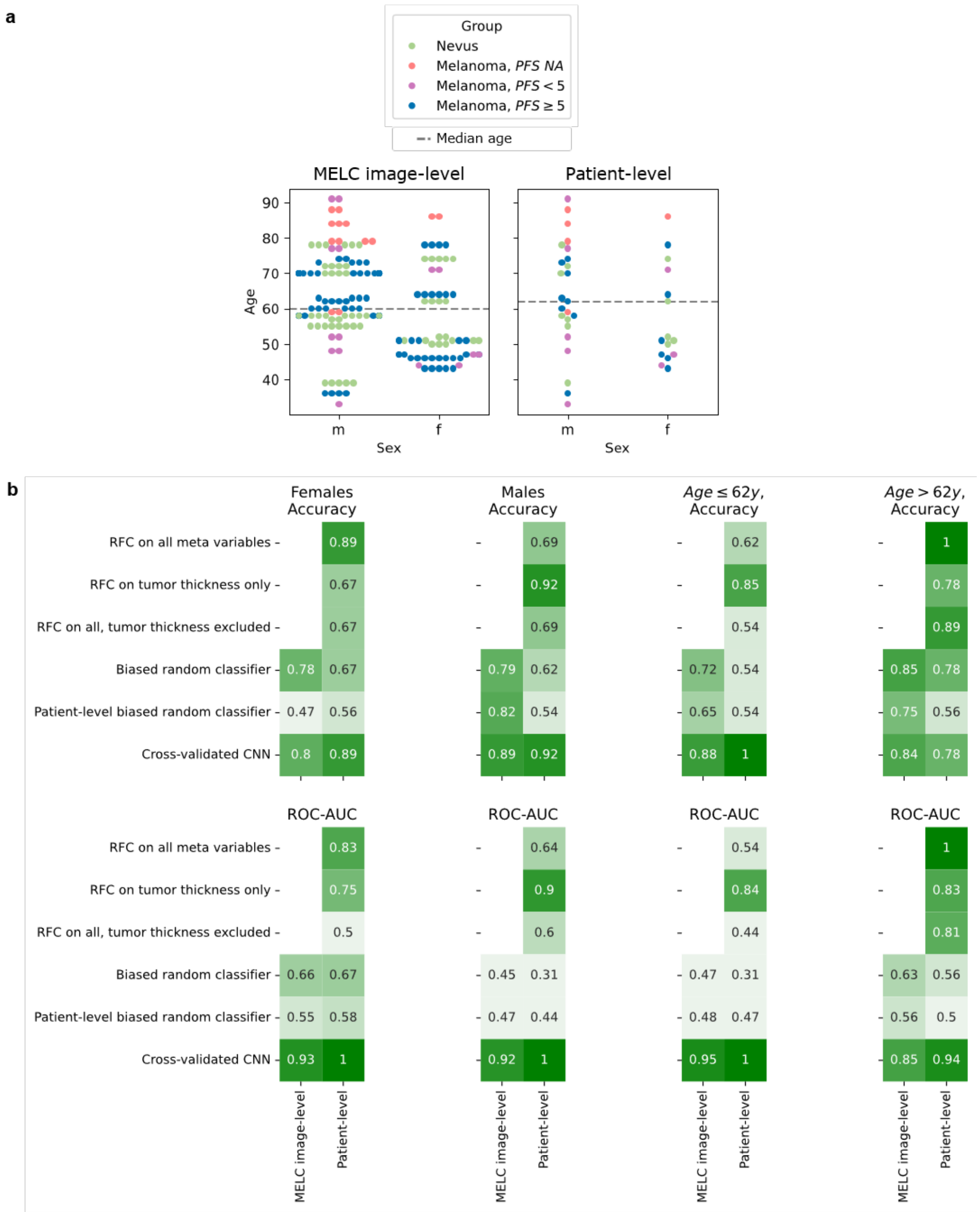


Figure S1: Results of sex- and age-resolved analyses. **a** Sex and age distributions in the groups. There are 24 men and 15 women in total (BN: 7m/5f; MM with PFS $=$ N/A: 4m/1f; MM with PFS $< 5$: 5m/3f; MM with PFS $\geq 5$: 8m/6f). The mean age is 61.59 years (BN: 59.83 years; MM with PFS $=$ N/A: 79.2 years; MM with PFS $< 5$: 57.875 years; MM with PFS $\geq 5$: 58.93 years). Median age is 62 years. **b** Sex-resolved (male versus female, first two columns) and age-resolved (above versus below median, last two columns) performance analysis for the tested PFS prediction models.

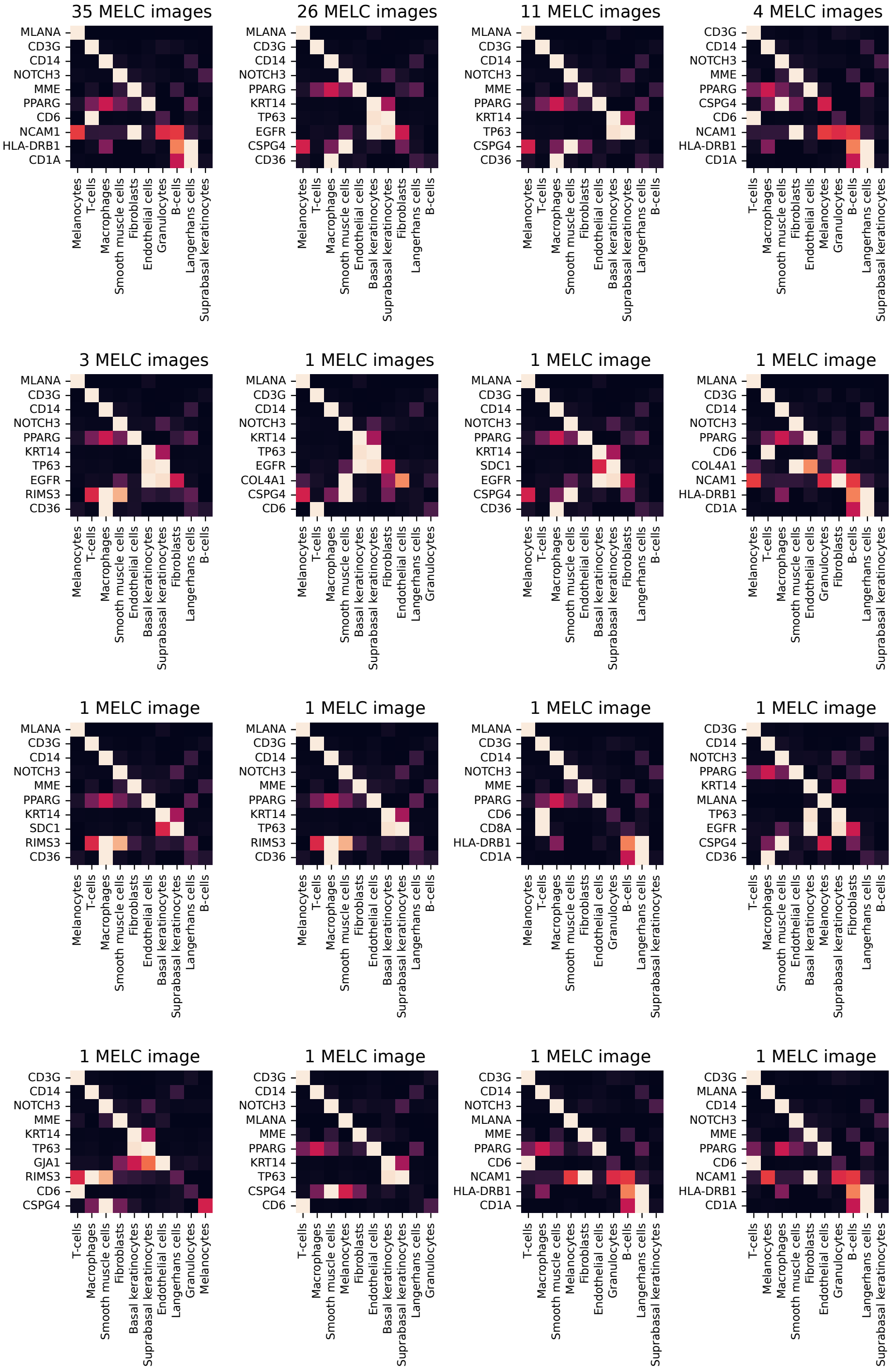


Figure S2: Cell type marker abundance pairs from HPA reference data used for our MELC data. Cell types and markers are sorted in the processing order employed by our iterative cell type assignment approach (cell types: left to right; markers: top to bottom). The plot titles indicate for how many MELC images the cell type-marker order displayed in the heatmaps has been used. For instance, the first heatmap visualizes that, for 35 MELC images, we first used MLANA abundance to identify melanocytes, then CD3G abundance to identify T-cells, and so on.

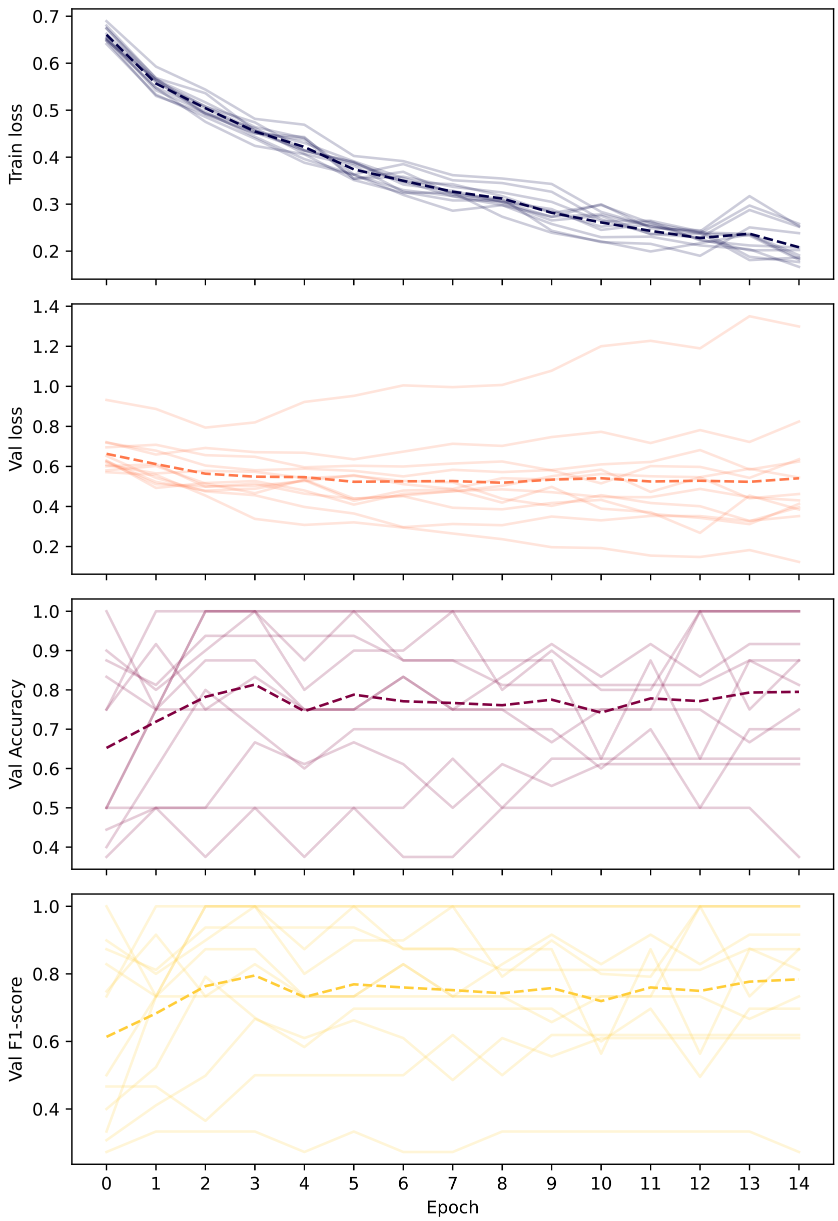


Figure S3: Development of training and validation loss, as well as validation accuracy and F1-score during the pre-training phase (BN vs. MM classification).

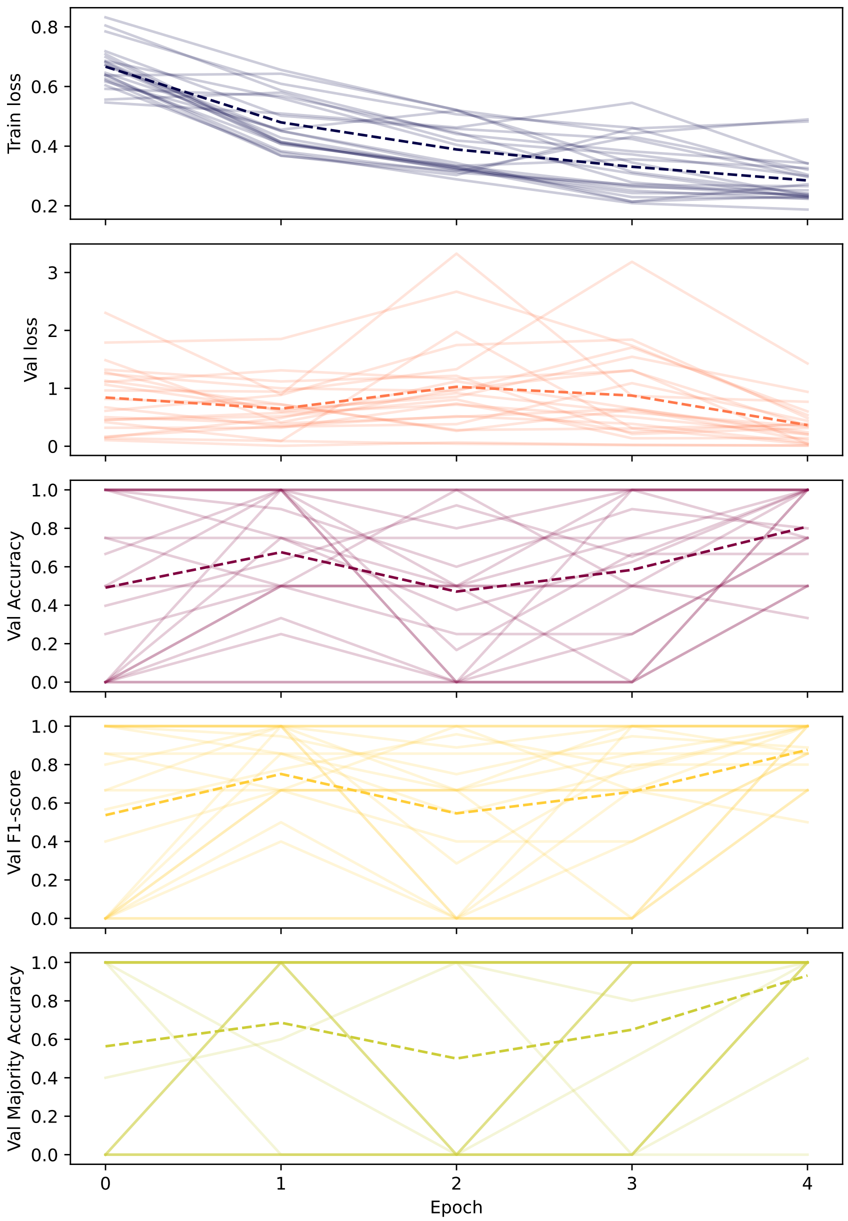


Figure S4: Development of training and validation loss, as well as validation accuracy and F1-score during the fine-tuning phase (PFS $< 5$ vs. PFS $\geq 5$ classification for MM samples).

#### 2 Supplementary Tables

Table S1: Full performance metrics of cross-validated approaches.

| | | MELC image-level | Patient-level |
|---|---|---|---|
| RFC on all meta variables | Accuracy | | 0.73 |
| | AUC-ROC | | 0.68 |
| | F1-score | | 0.57 |
| | Specificity | | 0.86 |
| | Sensitivity | | 0.50 |
| RFC on tumor thickness only | Accuracy | | 0.82 |
| | AUC-ROC | | 0.83 |
| | F1-score | | 0.78 |
| | Specificity | | 0.79 |
| | Sensitivity | | 0.88 |
| RFC on all except tumor thickness | Accuracy | | 0.64 |
| | AUC-ROC | | 0.56 |
| | F1-score | | 0.33 |
| | Specificity | | 0.86 |
| | Sensitivity | | 0.25 |
| Biased random classifier | Accuracy | 0.79 | 0.64 |
| | AUC-ROC | 0.54 | 0.42 |
| | F1-score | 0.13 | 0.00 |
| | Specificity | 0.87 | 1.00 |
| | Sensitivity | 0.13 | 0.00 |
| Patient-level biased random classifier | Accuracy | 0.70 | 0.55 |
| | AUC-ROC | 0.51 | 0.48 |
| | F1-score | 0.13 | 0.00 |
| | Specificity | 0.87 | 1.00 |
| | Sensitivity | 0.13 | 0.00 |
| Cross-validated CNN | Accuracy | 0.86 | 0.91 |
| | AUC-ROC | 0.91 | 0.99 |
| | F1-score | 0.58 | 0.88 |
| | Specificity | 0.88 | 0.93 |
| | Sensitivity | 0.73 | 0.88 |

Table S2: Full performance metrics of adapted workflow where pre-training is carried out without the patients from the test folds of the LOOCV scheme for four randomly selected patients.

| | | MELC image-level | Patient-level |
|---|---|---|---|
| Pre-training on all | Accuracy | 0.83 | 1.0 |
| | AUC-ROC | 0.97 | 1.0 |
| | F1-score | 0.75 | 1.0 |
| | Specificity | 0.88 | 1.0 |
| | Sensitivity | 0.75 | 1.0 |
| Pre-training without CV sample | Accuracy | 0.67 | 1.0 |
| | AUC-ROC | 0.81 | 1.0 |
| | F1-score | 0.50 | 1.0 |
| | Specificity | 0.75 | 1.0 |
| | Sensitivity | 0.50 | 1.0 |

Table S3: Detailed discussion of markers that are the most significantly differentially expressed within the cells of interest across the two patient cohorts ($PFS < 5$ and $PFS \geq 5$).

| Protein | Literature context |
|---|---|
| CD14 | The increased abundance of CD14 in PFS $< 5$ samples can directly be associated with the increased macrophage density observed for these samples. |
| CD4 | The role of CD4+ T cells in immunosurveillance of melanoma cells is highly discussed, and the full extent of their contribution is still unclear[1–3]. We observe a higher expression in primary tumors of patients with unfavourable outcome. |
| CDKN1B | Decreased nuclear CDKN1B expression has been supposed to correlate with greater tumor thickness, tumor progression, and poor survival[4–6]. In contrast to these studies, we observe a higher cell-level expression in the ROIs of the samples from patients with PFS $< 5$. |
| KRT14 | Many studies have reported progression-related differential KRT14 expression, but observations on the direction of the difference diverge in literature[7–10]. We observe a higher abundance in the primary tumors of the group with favorable outcome. |
| NCAM1 (CD56) | Despite the functional role of NCAM1 in MM still being unclear, a higher expression has been associated to higher tumor aggressiveness in melanoma[11,12]. Our data support this association. |
| TP73 | TP73 has been reported as reliant predictor for prognosis and progression in other cancer types such as cervical cancer[13] and has been suggested to influence the invasiveness of MM[14,15]. We observe a lower expression in the aggressive samples. |
